# Beyond rumination: an ecological momentary assessment of daily thinking patterns in maladaptive daydreaming

**DOI:** 10.64898/2026.10.29.762728

**Authors:** Annie S. Lu, Melissa Reich-Fuehrer, Eric S. Andrews, Sarah Hennessy, Kalina Christoff Hadjiilieva, Julia W. Y. Kam, Jessica R. Andrews-Hanna

**Affiliations:** Department of Psychology, University of Arizona, Tucson, Arizona, United States of America; Department of Applied Psychology, Bouvé College of Health Sciences, Northeastern University, Boston, Massachusetts, United States of America; Brain and Creativity Institute, University of Southern California, Los Angeles, California, United States of America; Department of Psychology, University of British Columbia, British Columbia, Canada; Department of Psychology, University of Calgary, Alberta, Canada; Hotchkiss Brain Institute, University of Calgary, Alberta, Canada; Cognitive Science Program, University of Arizona, Tucson, Arizona, United States of America

**Keywords:** Maladaptive daydreaming, maladaptive repetitive thoughts, rumination, mind-wandering, ecological momentary assessment (EMA), mental health, psychological well-being

## Abstract

Maladaptive daydreaming is a form of maladaptive repetitive thought characterized by excessive, dysfunctional fantasy engagement, with notable links to psychological distress and poor well-being. Although symptoms can be reliably measured along a continuum, it remains unclear how they manifest in the phenomenological characteristics of everyday thinking, as well as which thought characteristics are unique to maladaptive daydreaming versus shared with trait rumination, a transdiagnostic manifestation of maladaptive repetitive thinking that frequently co-occurs with maladaptive daydreaming. To advance these questions, we conducted a pre-registered study using ecological momentary assessment to track the everyday thoughts of 446 young adults (aged 18 to 34) spanning a range of maladaptive daydreaming symptom severity as assessed with the 16-item Maladaptive Daydreaming Scale. Higher maladaptive daydreaming scores were associated with several poor mental health indicators, including higher trait rumination. Positive associations between maladaptive daydreaming scores and both negatively-valenced and task-unrelated thinking emerged, yet these relationships were accounted for by variance shared with rumination. Beyond rumination, higher maladaptive daydreaming scores were uniquely associated with worse mood, as well as greater internally-oriented attention, more socially-oriented thought, and greater overall thought persistence. Maladaptive daydreaming also uniquely moderated the relationship between internal orientation and social orientation, such that the positive association between internal and social thought content became stronger at higher levels of maladaptive daydreaming. Overall, these findings provide novel evidence that maladaptive daydreaming levels are linked to unique socioemotional features and restricted dynamics, beyond its overlap with general ruminative distress in everyday life. These insights suggest that clinical interventions should target immersive fantasy engagement alongside transdiagnostic maladaptive repetitive thinking.

## Introduction

Maladaptive daydreaming (MD), first defined by Somer [1], describes a form of extensive fantasy activities that disrupts daily academic, interpersonal, and/or vocational functioning. Individuals with high levels of MD symptoms reported excessive and repetitive fantasizing, often accompanied by disengagement from reality [2–4], distress about losing control over their daydreaming [5], and reliving unsatisfactory past events to compensate for failures, unfulfilled interpersonal needs, or real-world distress [4]. While a growing number of studies have examined MD in online social media communities and among clinical populations, relatively little is known about how MD symptoms manifest in everyday cognitive experiences.

Clinically, while MD is not recognized in standard mental health diagnosis, empirical studies have viewed it as a valid and reliable construct [6,7]. Characterized as addictive-compulsive immersion in vivid fantasy [8], MD shares comorbidity and empirical links with other mood disorders and clinical conditions such as depression, obsessive-compulsive disorder (OCD) [9], attention-deficit/hyperactivity disorder (ADHD) [10], dissociative disorders [8], and behavioral addiction [5].

Theoretical work [11] positions MD within the Dynamic Framework of Thought [12] by suggesting that the phenomenon’s compulsive and habitual features are more accurately characterized as compulsive fantasy rather than “maladaptive daydreaming.” More specifically, Burrell and colleagues propose a profile of restricted dynamics in compulsive fantasy emerging from heightened automatic constraints on thought, combined with intermittent intentional allocation of attentional resources (called deliberate constraints) [11,12]. Although we retain the term ‘maladaptive daydreaming’ (or MD) throughout this manuscript to align with measurement scales, our empirical findings aim to understand the broader everyday thinking patterns of individuals who more strongly endorse maladaptive compulsive fantasizing.

Cognitively, some studies have highlighted that MD symptoms may share similarities with forms of maladaptive repetitive thinking spanning rumination [13], obsessive-compulsive thinking, and recurring trauma-related compensatory episodes [9]. Trait rumination – a well-established form of maladaptive repetitive thinking characterized by persistent, involuntary, and negatively-valenced thoughts [14–18] – moderately correlates with MD in prior research (*r* = .49) [19]. MD is also often associated with depressive symptoms, for which rumination is a central feature [17]. However, the extent to which MD reflects the similar cognitive processes as rumination remains unclear.

Establishing whether MD represents a phenomenologically distinct construct or merely a variant of rumination carries critical theoretical and clinical implications. While episodes of compulsive fantasy in MD may offer short-term relief or reward, the experience has been associated with longer-term maladaptive outcomes [20]. Beyond the temporary affective qualities that may distinguish episodes of MD from state rumination, the social themes described in MD-related thoughts may also differentiate the experience from rumination, as rumination tends to be conceptualized as a more self-focused phenomenon [15]. Comparing MD and trait rumination using an ecological momentary assessment (EMA) approach is therefore essential to clarify their shared and distinct characteristics.

While a growing body of research has been devoted to characterizing the nature of acute MD experiences (e.g. the process of engaging in maladaptive daydreaming), the daily thinking patterns of individuals with varying levels of MD remain under-explored. The EMA approach has been widely employed in research on daydreaming and mind-wandering, enabling the sampling of experiences outside laboratory settings and the capture of real-time thoughts, affect, and behavior [21,22]. However, to our knowledge, EMA has not yet been applied to examining relationship between MD and broader characteristics of everyday thought, while accounting for trait rumination. Existing related work capturing temporary characteristics of MD included a daily diary study, which demonstrated how obsessive-compulsive disorder symptoms, dissociation, and negative emotions are accompanied by MD [23]. Another daily diary study identified associations between sleep disturbance and mind wandering, though not MD severity [24]. While these findings offer valuable insight into the longitudinal features of MD, many questions about the MD symptoms remain poorly understood, likely due to the relative new construct and its limited recognition as a distinct phenomenon. Specifically, unresolved questions include: (1) how trait MD manifests in the real-time characteristics of everyday thoughts, (2) which momentary cognitive characteristics are uniquely associated with MD versus shared with general transdiagnostic processes like trait rumination; and (3) whether trait MD moderates the dynamics between different dimension of daily thought characteristics.

### The Present Study

The present study addresses existing research gaps by using EMA data to enable real-time assessments of everyday thoughts from individuals with different levels of MD. We utilized EMA data collected via a cross-platform smartphone app – called Mind Window [25] – to examine associations between MD and daily thought characteristics. This preregistered study aimed to characterize the daily cognitive style across different MD levels by examining its unique and overlapping features relative to trait rumination. Specifically, we tested which momentary thought characteristics were uniquely associated with MD symptoms and which were shared with trait rumination, thereby clarifying the distinct role of MD in shaping mental health, well-being, and daily cognitive patterns.

#### Preregistered Hypotheses

We tested a set of mental health indices in relation to MD to replicate previously reported relationships. We hypothesized that higher levels of MD would be positively associated with higher levels of rumination [19]. Consistent with prior work linking MD symptoms to psychological distress and psychopathology [10,19,26–28], we hypothesized that MD scores would be positively associated with loneliness, anxiety symptoms, and depressive symptoms, and negatively associated with trait mindfulness and well-being.

We next examined if MD symptoms was associated with daily thought characteristics measured by EMA. Given the imagery-intensive nature of MD episodes, we hypothesized that higher MD scores would be linked to a greater degree of internally oriented attention (internal-orientation of thoughts, such as memories, imagination, etc.), but a lower degree of external-orientation (for example, outside environment) in daily experience. Although MD has been reported to temporarily enhance mood due to possible compensatory contents [7], its overall association with psychological distress [23,27,28] led us to hypothesize that higher MD scores would be associated with more negative state affect and more negative thought content in general. Furthermore, because MD has been proposed to substitute for real-world social experiences and involve an imagined ideal self as well as idealized social interactions [2,4,29,30], we hypothesized that higher MD scores would be associated with higher degrees of self- and social-orientation during internally-oriented mental activity.

Given MD’s characteristic tendency towards sticky, unprompted, internally generated, and task-unrelated thoughts, we tested a preregistered set of moderation hypotheses examining whether MD shape relationships among thought dynamics including intentionality, task-relatedness, persistence, and related dimensions. Full details of preregistered hypotheses are available in the preregistration (https://osf.io/86quy).

## Methods

The Institutional Review Board responsible for human subjects research at The University of Arizona reviewed this research project and found it to be acceptable, according to applicable state and federal regulations and University policies designed to protect the rights and welfare of participants in research.

### Data Collection and Cleaning

Participants enrolled for the study through the SONA system at the University of Arizona during the Fall 2021, Spring 2022, and Fall 2022 semesters. Data were accessed Oct 24, 2024 for research purposes. The research team had access to information that could identify individual participants (email), while identifiable information was stored separately from the primary survey and EMA dataset in a restricted Box Health folder. All datasets were fully anonymized prior to analyses.

Participants were required to be fluent in English and at least 18 years old. Following online informed consent, participants initially completed the MDS-16 questionnaire in Qualtrics [7], as well as a number of other questionnaires not analyzed in the present study. Participants received credit for completion of the study through the SONA system, as part of their course research requirements.

As part of the study, participants downloaded and used the cross-platform Mind Window smartphone app (version 1.5.6) [25] to provide EMA data, which was developed to build a comprehensive database of everyday thoughts and identify cognitive profiles both quantitatively and qualitatively. Upon registration, participants anonymously provided their demographic information and completed a set of selected questionnaires, including mental health indices such as Patient-Reported Outcomes Measurement Information System (PROMIS) [31], trait rumination [15], mindfulness [32], and other well-being measures. After completing the required questionnaires and supplying their waking hours, participants received 6 push notifications during the waking hours, prompting them to complete the EMA survey. Every notification was valid for 10 minutes, after which the survey became unavailable until the next prompt. Each EMA survey was composed of 12 core questions (in random order), 2 additional questions from a rotating question pool, and a free response section. These EMA surveys were measured repeatedly to quantitatively and qualitatively capture the momentary mental activity of the individual.

### Participant Sample

A total of 719 participants contributed data to the questionnaires and EMA protocol. Of these, 446 participants were included in the present analyses because they had both a completed MDS score and at least 10 EMA check-ins. The remaining 273 participants were excluded because they did not have a completed MDS score. 446 participants were included with ages ranging from 18 to 34 (M = 19.34, SD = 2.44). See Table 1 for a summary of demographic information. 14,351 completed EMA surveys were collected across participants. Participants provided a median of 31 EMA check-ins (range = 10–336) across a median of 9 calendar days with at least one completed check-in (range = 3–176). On these days, participants completed a median of 3.36 check-ins per day, or approximately 56% of the six check-ins per day.

**Table 1.** Demographic characteristics of the study sample (N = 446).

| **Characteristic** | **N (%)** |
|---|---|
| Age (Mean, SD) | 19.3 (2.44) |
| **Gender** |  |
| Cis Woman | 356 (79.8%) |
| Cis Man | 80 (17.9%) |
| Gender Minority | 10 (2.2%) |
| **Education** |  |
| Less than High School | 1 (0.2%) |
| High School or GED | 170 (38.1%) |
| Some College, College Degree, or Higher | 275 (61.7%) |
| **Employment** |  |
| Full-Time | 32 (7.2%) |
| Other | 3 (0.7%) |
| Part-Time | 119 (26.7%) |
| Student | 199 (44.6%) |
| Unemployed | 93 (20.9%) |
| **Race/Ethnicity** |  |
| Non-Hispanic White | 227 (50.9%) |
| Latinx, Hispanic, or Hispanic American | 117 (26.2%) |
| Asian or Asian American | 44 (9.9%) |
| Multiracial | 27 (6.1%) |
| Black or African American | 20 (4.5 %) |
| Native American, Alaska Native, First Nations, or Indigenous | 4 ( 0.9%) |
| Arab/Middle Eastern | 1 (0.2%) |
| Native Hawaiian or Other Pacific Islander | 1 (0.2 %) |
| Prefer not to answer | 5 (1.1%) |

Values are presented as mean (SD) for age and N (%) for other information.

### Trait Questionnaires

A series of validated questionnaires were administered to assess traits and well-being. All questionnaires were scored according to their scoring procedures.

#### Maladaptive Daydreaming

MD level was measured as participants’ scores on the 16-item Maladaptive Daydreaming Scale (MDS-16) [7,33]. The scale was designed to measure the severity of MD symptoms [7,33], including subscales of yearning, impairment, kinesthetic movement, as well as two additional items measuring initiation and maintenance of music. Because this study focuses on the extent to which MD interferes with daily thought characteristics, our primary analyses emphasized overall MD severity as measured by the full scale. We also examined the individual subscales as preregistered and explored their respective associations with trait rumination.

Participants answered each question on a 10-point scale. Higher scores reflect greater MD symptoms. It showed excellent internal consistency in the present sample, Cronbach’s α = .94, 95% CI [0.93, 0.95].

#### Rumination

Trait rumination was measured using the 12-item rumination subscale from the Rumination-Reflection Questionnaire (RRQ) [15]. Participants answered each question using a 5-point scale from *Strongly Disagree (1)* to *Strongly Agree (5)*. The subscale includes items such as “I tend to ruminate or dwell over things that have happened to me for a long time afterwards.” Higher scores reflect greater tendency to engage in ruminative thinking. It showed excellent internal consistency in the present sample, Cronbach’s α = .91, 95% CI [0.90, 0.93].

#### Mindfulness

Trait mindfulness was measured with the 15-item Mindful Attention Awareness Scale (MAAS) [32]. Participants answered each question using a 6-point scale from *Almost always (1)* to *Almost Never (6)*. An example item includes “I do jobs or tasks automatically, without being aware of what I’m doing.” Higher scores reflect a greater tendency to stay aware and nonjudgmental of the present moment. It showed excellent internal consistency in the present sample, Cronbach’s α = .91, 95% CI [0.90, 0.92].

#### Well-being: positive relation with others

Positive relation with others was measured through a 3-item subscale of the Ryff’s Psychological Well-being Scale using a 7-point scale from *Strongly Agree (1)* to *Strongly Disagree (7)* [34]. Higher scores reflect greater interpersonal well-being. Positive relation with others showed questionable internal consistency in the present sample, Cronbach’s α = .60, 95% CI [0.53, 0.66. Thus, the analysis should be interpreted with caution.

#### Loneliness

Loneliness was measured through the Three-Item Loneliness Scale (T-ILS) using a 3-point scale from *Hardly Ever* (1) to *Often (3)* [35]. An example item includes “I feel isolated from others”. Higher scores reflect high loneliness levels. It showed good internal consistency in the present sample, Cronbach’s α = .85, 95% CI [0.82, 0.87].

#### Anxiety

Anxiety level was measured using the 4-item Patient-Reported Outcomes Measurement Information System - Anxiety (PROMIS-A) using a 5-point scale from *Never (1)* to *Always (5)* [31]. Higher scores reflect greater severity of anxiety symptoms. It showed good internal consistency in the present sample, Cronbach’s α = .88, 95% CI [0.86, 0.90].

#### Depressive symptoms

Depression level was measured using the 4-item Patient-Reported Outcomes Measurement Information System - Depression (PROMIS-D) using a 5-point scale from *Never (1)* to *Always (5)* [31]. Higher scores reflect greater severity of depressive symptoms. It showed excellent internal consistency in the present sample, Cronbach’s α = .94, 95% CI [0.93, 0.95].

### EMA Measures

EMA measures included a set of questions about daily thought characteristics. Internal-orientation (participants reported that their momentary attention was oriented towards thoughts/memories), external-orientation (momentary attention oriented towards the external environment), and interoceptive-orientation (bodily sensations) together captured a thought’s attentional orientation. Participants’ state affect valence captured to what extent their momentary feelings were positive or negative, while affective content measured the valence of the emotional content of the momentary thought itself. Degrees of self- and social-orientation described the degree to which each thought was about one’s self and/or other(s). Persistence described how long the thought was expected to be if not interrupted by notification. Task-relatedness described the degree to which thoughts/attention was focused on the primary “task” or “activity” at hand. Intentionality described the degree to which participant actively direct their thought. Participants also indicated who they were with at the time of the survey as a social context. The full list of EMA measures is available in supplemental materials (**S1. EMA measures**).

### Statistical Analysis

Several EMA variables were recoded. For the measurement of internally-oriented thoughts, we defined a thought to be internally-oriented if the degree of internal-orientation rating exceeded 0.5 (out of 1). Based on this definition, for analyses that were specifically interested in internally-oriented thoughts, we classified a subset of EMA observations which were composed of solely internally-oriented thoughts. This subset was further restricted to participants who contributed more than five such internally-oriented thought observations. For the measurement of social context, we dummy-coded a dichotomous variable to index being alone. Responses were coded as 1 (alone) if participants selected “No one/alone,” “a pet,” or both, and 0 (not alone) if any other social categories were endorsed.

MD was treated as a continuous measure to preserve variability in levels and maximize statistical power. To assess whether the relationship between MD levels and the thought characteristic of interest was independent of general rumination tendencies, rumination was added as a covariate to test whether the MD–outcome association remained significant after adjusting for shared variance with trait rumination. We included gender as a covariate in models examining mental health–related variables [36], and we included age as a covariate in models involving self-focused thinking, considering an age effect in self-referential processes [37].

Statistical analyses were performed in R 4.4.1 [38]. To determine the relationship between MD and other thought characteristics, linear regression models and multilevel models were used to examine the relationship with MD level being a predictor variable or a moderator. Model assumptions were assessed using visual inspection of diagnostic plots and variance inflation factors (VIFs) to examine linearity, heteroscedasticity, normality, and collinearity.

Linear regression analyses were conducted using the lm() function from the base stats package. We applied lmrob() from the robustbase package [39] in case of violation of assumptions. All predictor variables were mean-centered. Detailed linear regression tables are provided in the Supplementary Materials (**S2. Linear Regression Analyses**). Considering the nested data structure (repeated measurements clustered within participant), we adopted multilevel modeling when assessing the link between MD (between-subject variable) and EMA sampled variables of interest (within-subject variable). Multilevel models were estimated using the lmer() function from the lmerTest package [40] with random intercept of subject ID to control for within-subject dependency. We estimated robust standard errors using the clubSandwich package [41] in case of violation of assumptions. Robust standard errors were computed using the CR2 estimator (Cluster-Robust Variance Estimator, type 2), which provides a small-sample corrected sandwich estimator that adjusts for within-cluster dependence and heteroscedasticity. This method was implemented using the vcovCR() function, with hypothesis tests conducted using the coef_test() function and Satterthwaite-approximated degrees of freedom. Predictors were grand-mean centered if they were level-2 predictors, and predictors were within-person-mean-centered if they were level-1 momentary predictors. Missing individual EMA prompts were omitted at the observation level without dropping participants from Level-2 analyses.

We report standardized coefficients alongside unstandardized coefficients for completeness. Standardized coefficients were obtained by refitting each model using standardized continuous variables. For multilevel models, standardized variables were entered into the same mixed-effects model specification as the primary analyses, including the same random-effects structure, and thus standardized coefficients (*β*) for continuous predictors represent the expected change in standard deviation units.

In the current study, statistical significance was evaluated using a conventional alpha level of 0.05. To control for the inflation of false positives across our multiple mental health relationships and EMA thought characteristics, p-values were adjusted (p_adj_) using the Benjamini-Hochberg False Discovery Rate (FDR) procedure across hypotheses.

## Results

### Statistical Reporting Approach

To evaluate whether the relationships between MD and our outcomes of interest were independent of general repetitive negative thinking tendencies, all primary linear and multilevel models were evaluated both with and without trait rumination included as a covariate. Unless explicitly stated otherwise, all significant main effects and interactions of MD reported below remained statistically significant (*p* < 0.05) after controlling for trait rumination. Cases where rumination attenuated or modified the MD effect are noted explicitly within their respective subsections. Full model statistics for all unadjusted and adjusted models are provided in Tables 2–5 and Supplementary Materials S2–S4.

**Table 2.** Associations between Maladaptive Daydreaming and Mental Health Outcomes Controlling for Rumination. Unstandardized coefficients (B), 95% confidence intervals (CI), standard errors (SE), p-values, and standardized coefficient (β) are presented for linear regression models examining the association between MD and rumination, mindfulness, interpersonal well-being (positive relations with others), loneliness, anxiety, and depression.

| *Outcome* | *B (MD)* | *95% CI* | *SE* | *β* | *p* |
|---|---|---|---|---|---|
| Mindfulness | -0.0148 | (-0.0198–-0.0099) | 0.0025 | -0.295 | <0.001 |
| Interpersonal Well-being | -0.0533 | (-0.0728–-0.0339) | 0.0099 | -0.260 | <0.001 |
| Loneliness | 0.0192 | (0.0095–0.0290) | 0.0050 | 0.191 | <0.001 |
| Anxiety | 0.0591 | (0.0418–0.0763) | 0.0088 | 0.276 | <0.001 |
| Depression | 0.0670 | (0.0438–0.0902) | 0.0118 | 0.276 | <0.001 |

**Table 3.** Multi-level Relationship between Maladaptive Daydreaming and Internal-orientation Controlling for Trait Rumination.

| *Predictors* | *Estimates (95% CI)* | *SE* | *t-stats* | *p* |
|---|---|---|---|---|
| (Intercept) | 0.4245 (0.4112–0.4378) | 0.0068 | 62.778 | <0.001 |
| MD | 0.0011 (0.0003–0.0018) | 0.0004 | 2.795 | **0.005** |
| Rumination | 0.0012 (-0.0168–0.0191) | 0.0091 | 0.130 | 0.897 |
| σ<sup>2</sup> | 0.06 |  |  |  |
| τ<sub>00 ID</sub> | 0.02 |  |  |  |
| ICC | 0.25 |  |  |  |
| N<sub>ID</sub> | 446 |  |  |  |
| Observations | 14165 |  |  |  |
| Marginal R<sup>2</sup> / Conditional R<sup>2</sup> | 0.005 / 0.224 |  |  |  |

Estimates are unstandardized coefficients (B) with 95% CI, standard errors (SE), test statistics, and *p*-values. Significant results are marked in bold. Random effects include random intercepts for participants (ID). Model fit indices include residual variance (σ²), intercept variance (τ₀₀), ICC, and marginal and conditional R².

**Table 4.** Multi-level Relationship Between Maladaptive Daydreaming and Self-Reported State Affect (Internal Thoughts Only).

|  | (A) Outcome: state affect |  |  |  | (B) Outcome: state affect (controlling for rumination) |  |  |  |
|---|---|---|---|---|---|---|---|---|
| *Predictors* | *B (95% CI)* | *SE* | *Statistic* | *p* | *B (95% CI)* | *SE* | *Statistic* | *p* |
| (Intercept) | 0.6224 (0.6065–0.6383) | 0.0081 | 76.889 | <0.001 | 0.6241 (0.6085–0.6397) | 0.0079 | 78.541 | <0.001 |
| MD | -0.0015 (-0.0024–-0.0005) | 0.0005 | -3.056 | **0.003** | -0.0009 (-0.0018–0.0000) | 0.0005 | -1.892 | 0.061 |
| Rumination | / | / | / | / | -0.0507 (-0.0741–-0.0273) | 0.0118 | -4.292 | **<0.001** |
| σ<sup>2</sup> | 0.06 |  |  |  | 0.06 |  |  |  |
| τ<sub>00 ID</sub> | 0.02 |  |  |  | 0.02 |  |  |  |
| ICC | 0.25 |  |  |  | 0.25 |  |  |  |
| N<sub>ID</sub> | 370 |  |  |  | 370 |  |  |  |
| Observations | 5276 |  |  |  | 5276 |  |  |  |
| Marginal R<sup>2</sup> / Conditional R<sup>2</sup> | 0.010 / 0.253 |  |  |  | 0.026 / 0.254 |  |  |  |

(A) model without rumination. (B) model with rumination. Estimates are unstandardized coefficients (B) with 95% CI, standard errors (SE), test statistics, and p-values. Significant results are marked in bold. Random effects include random intercepts for participants (ID). Model fit indices include residual variance (σ²), intercept variance (τ₀₀), ICC, and marginal and conditional R².

**Table 5.** Summary of Additional Preregistered Analyses.

| **Hypothesis** | **Outcome** | **B** | **SE** | **p** | **β** | **Significant after controlling for rumination?** |
|---|---|---|---|---|---|---|
| Higher MD → lower intentionality | Intentionality | -0.0004 | 0.0004 | 0.335 | -0.02 | No |
| MD moderates intentionality → persistence | Interaction | -0.0006 | 0.0006 | 0.355 | -0.01 | No |
| MD moderates internal-orientation → persistence | Interaction | -0.0001 | 0.0006 | 0.812 | 0.00 | No |
| MD moderates external-orientation → persistence | Interaction | 0.0004 | 0.0006 | 0.461 | 0.01 | No |
| MD moderates intentionality → social-orientation | Interaction | -0.0006 | 0.0008 | 0.503 | -0.01 | No |
| MD yearning moderates positive content → persistence | Interaction | 0.0007 | 0.0004 | 0.127 | 0.01 | No |
| MD impairment moderates task-relatedness → persistence | Interaction | -0.0010 | 0.0007 | 0.125 | -0.02 | No |

*Note. B* = unstandardized regression coefficient; *SE* = standard error; *β* = standardized regression coefficient. Statistical significance was evaluated at p < 0.05. Interaction terms represent moderation effects.

### Descriptive Statistics

Across the sample, participants’ MDS-16 scores ranged from 0.00 - 89.38, with a mean of 25.92 across participants (median = 23.12; SD = 18.65), and a positive skew (skewness = 0.77) (Fig 1). Robust statistical approaches were employed where appropriate to reduce the influence of skewness. Primary analyses treated MD as a continuous measure (see Methods); 92 of these participants (20.6%) exceeded the commonly-used clinical screening cutoff (40 in MDS) of MD (English version; [7,42,43]).

**Fig 1.**
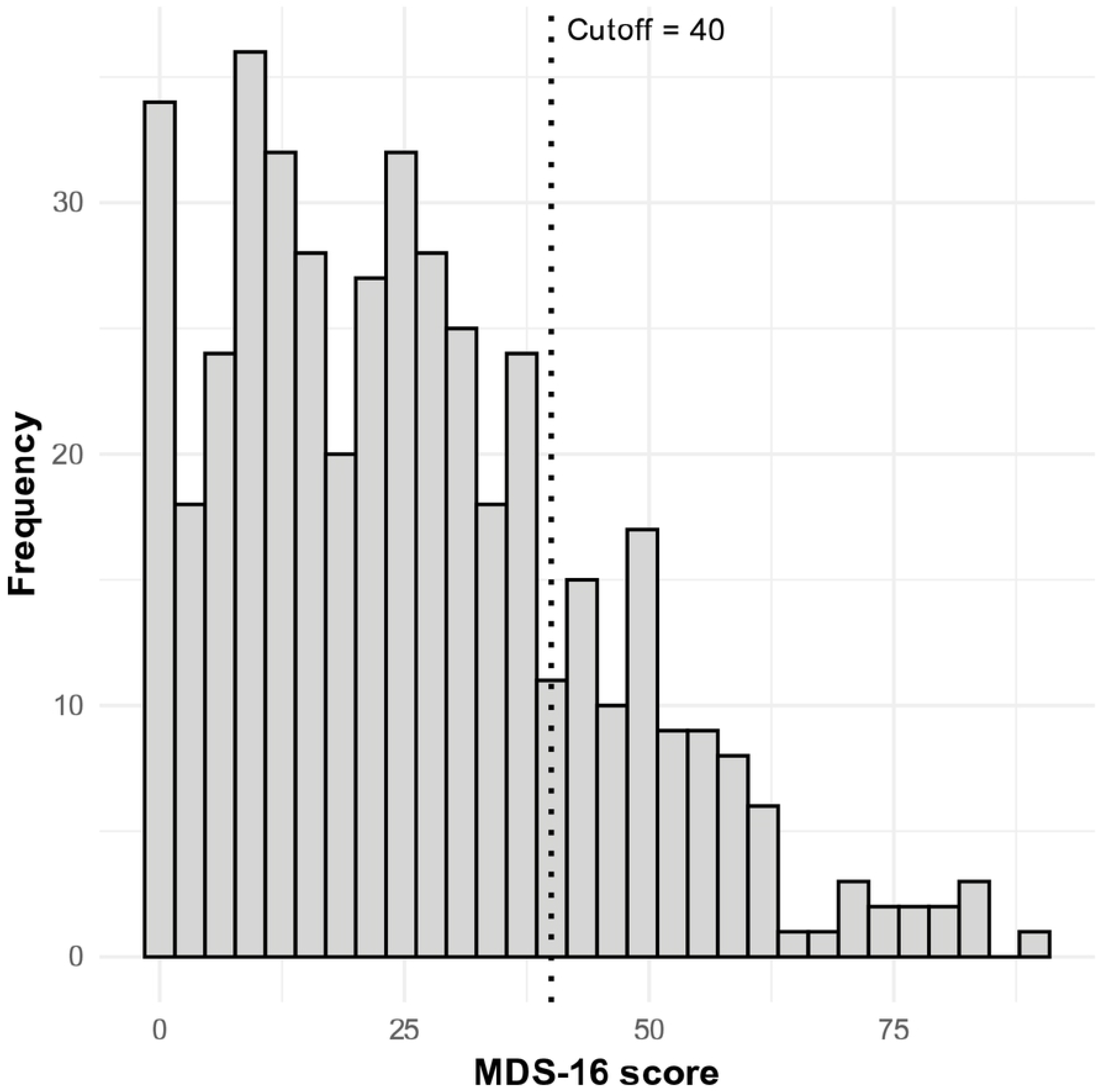
Distribution of MDS-16 scores. Histogram plot of Maladaptive Daydreaming Scale (MDS-16) total scores (N = 446). Scores range from 0 to 89.38, with higher scores reflecting greater MD. The vertical dotted line represents a screening cutoff score of 40 used in the literature [3].

### Relationship with Mental Health Indices

Consistent with our preregistered hypotheses, higher MD scores were associated with greater trait rumination, *B* = 0.0108, *SE* = 0.0018, *β* = 0.26, *p* < 0.001 (Fig 2A). Without rumination as a covariate, Higher MD scores were linked to lower trait mindfulness, *B* = -0.0214, *SE* = 0.0024, *β* = -0.43, *p* < 0.001 (Fig 2B). Higher MD scores were also associated with lower interpersonal well-being, *B* = -0.0609, *SE* = 0.0098, *β* = -0.30, *p* < 0.001 (Fig 2C), and greater loneliness, *B* = 0.0291, *SE* = 0.0049, *β* = 0.29, *p* < 0.001 (Fig 2D). In addition, MD scores were positively associated with anxiety, *B* = 0.0796, *SE* = 0.0091, *β* = 0.37, *p* < 0.001 (Fig 2E), and depressive symptoms, *B* = 0.0807, *SE* = 0.0099, *β* = 0.37, *p <* 0.001 (Fig 2F). Supporting our preregistered hypotheses, the MD impairment subscale was positively associated with both anxiety (*B* = 0.0796, *SE* = 0.0091, *β* = 0.38, *p* < 0.001) and depression (*B* = 0.0807, *SE* = 0.0099, *β* = 0.38, *p* < 0.001).

**Fig 2.**
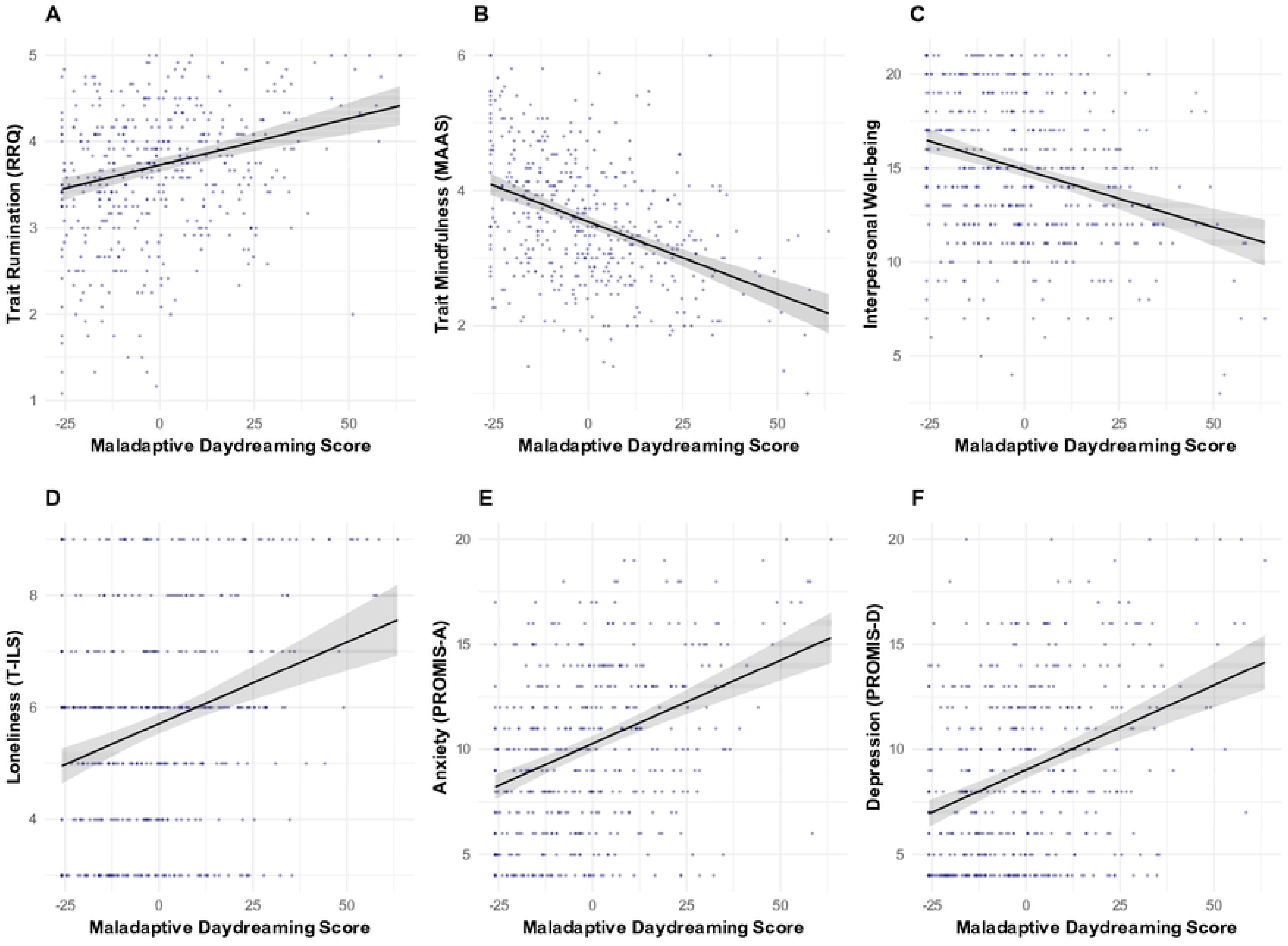
Linear regression between MD scores and mental health indices. (A) trait rumination; (B) trait mindfulness; (C) positive relation with others, (D) loneliness, (E) anxiety, and (F) depressive symptoms. MD scores were grand mean centered. The black line indicates the beta coefficient with 95% CI in gray shading. Each point represents an individual response.

As shown in Table 2, higher MD scores were uniquely associated – beyond variance shared with trait rumination – with lower trait mindfulness, lower interpersonal well-being, greater loneliness, and elevated anxiety and depressive symptoms.

### Multi-level Modeling Relationships with EMA-derived Momentary Thought Characteristics

#### MD was related to more inward-focused thought

We examined the relationship between MD scores and momentary attentional orientation, specifically the degree to which participants’ were engaged in internal thoughts (internally-oriented), external sensations (externally-oriented) and/or bodily sensations (interoceptive) (Table S3.1 – S3.3). In the unconditional model of internally-oriented attention, the ICC was 0.22, indicating that approximately 22% of the variance in momentary internal-orientation was attributable to between-person differences. Confirming our preregistered hypothesis, MD was significantly associated with a higher degree of momentary internal-orientation, *B* = 0.0011, *SE* = 0.0004, *p* = 0.003, *β* = 0.07; Table 3 presents full model parameters including the trait rumination covariate. Conversely, MD was negatively associated with degree of external-orientation, *B* = -0.0011, *SE* = 0.0003, *p* = 0.005, *β* = -0.07. The association between MD and interoceptive-orientation was not statistically significant, *B* = -0.0001, S*E* = 0.0002, *p* = 0.759, *β* = -0.00. Trait rumination did not have a significant association with either thought orientation.

#### MD was uniquely related to negative state affect, but not affective thought content

Confirming our preregistered hypothesis, MD scores were negatively associated with state affect valence across all EMA data, *B* = -0.0016, *SE* = 0.0004, *p* < 0.001, *β* = -0.11, such that individuals with higher MD scores rated their momentary feelings as less positive / more negative. Trait rumination was also independently linked to lower state affect (Table S3.4).

We also examined this relationship specifically within the subset of surveys that participants endorsed as pertaining to internal thoughts as opposed to external perceptions or bodily sensations (surveys rated more than 50% internally-oriented). In the internally-oriented thought subset, the negative association between MD and state affect was statistically significant only in the model excluding trait rumination. When rumination was included in the model, the effect of MD on state affect attenuated to non-significance, as shown in Table 4. This pattern suggests that shared variance with rumination may explain MD’s relationship with negative feelings during internally-oriented thinking.

An exploratory analysis was also conducted to examined the relationship between MD scores and affective content of momentary thoughts (i.e., the emotional tone of the thought content itself). Across all EMA observations, MD was initially associated with more negative thought content, *B* = -0.0011, *SE* = 0.0003, *p* = 0.001, *β* = -0.08. However, this association was no longer significant after controlling for rumination (*B* = -0.0006, *SE* = 0.0003, *p* = 0.049, *p_adj_*= 0.076; Table S3.6). Among internally-oriented thoughts specifically, MD was associated with more negative thought content, *B* = -0.0011, *SE* = 0.0004, *p* = 0.006, *β* = -0.08; adding trait rumination attenuated the effect of MD to non-significance, *B* = -0.0006, *SE* = 0.0004, *p* = 0.115, *β* = -0.05 (Table S3.7). Together, these findings suggest that the association between MD and the emotional tone of thought content is largely accounted for by variance shared with trait rumination.

#### MD interacts with trait rumination to shape state affect

Given this result, we tested a cross-level three-way interaction between MD score, trait rumination, and internal-orientation predicting state affect (valence of momentary feelings). The interaction was significant, *B* = -0.0025, *SE* = 0.0008, *p* = 0.005, *β* = -0.03 (Table S3.8; Fig 3), indicating that the effect of MD on the relationship between the degree of internal-orientation and state affect differed depending on participants’ ruminative tendencies. For participants exhibiting low trait rumination, higher MD scores manifested as positive relationships between internal orientation and concurrent state affect, such that thoughts rated as more internally-oriented associated with more positive concurrent state affect. For those with lower MD scores and low trait rumination, thoughts rated as more internally-oriented associated with less positive (more negative) concurrent state affect. In contrast, opposite relationships were found for individuals with high trait rumination: as MD scores increased, relationships between internal orientation and state affect became more negative. In summary, the cross-way three-level interaction revealed that whether internal-orientation was experienced as positive or negative depended on the combination of MD and trait rumination: higher MD scores were associated with more positive internally-oriented thoughts only when trait rumination was low, whereas higher trait rumination was associated with increasingly negative internally-oriented thoughts, particularly among individuals with high MD scores.

**Fig 3.**
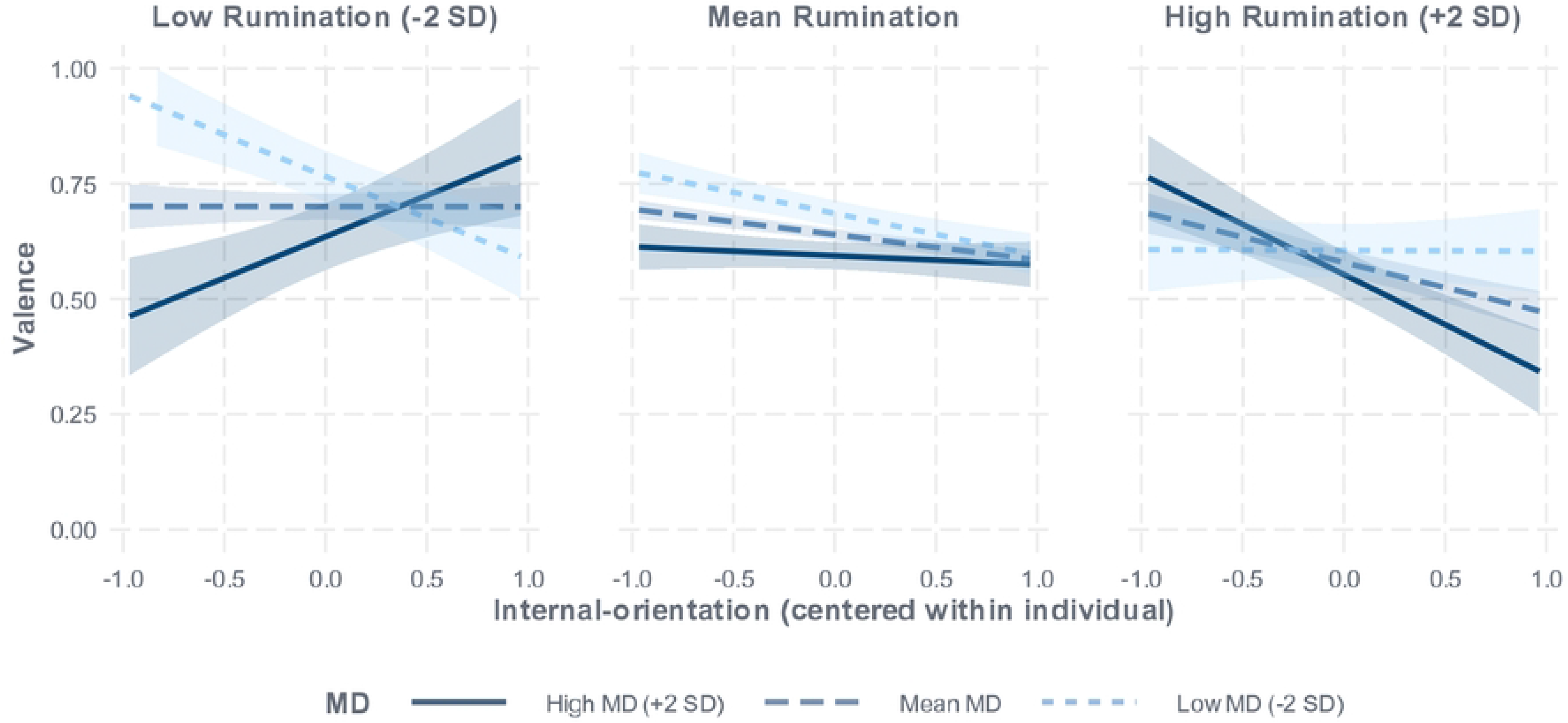
Three-way interaction between internal-orientation, MD, and trait rumination predicting valence. Simple slopes illustrate the association between internal-orientation and valence at low (−2 SD, light blue dotted line), mean (dark blue dashed line), and high (+2 SD, solid line) levels of MD for low (−2 SD, left panel), mean (middle panel), and high (+2 SD, right panel) levels of rumination. Lines indicate fitted regression slopes of the fixed effects, with shaded areas representing 95% confidence intervals. Internal-orientation ratings were centered within person. Simple slopes were probed at ±2 SD of the moderators to illustrate conditional effects across a wider range of observed values; all probing points fell within the observed data range. Because higher MD scores were sparsely represented due to skewness (N = 38 above +1.5 SD), estimates at the upper end of the MD distribution should be interpreted cautiously.

#### Higher MD scores predicted a higher degree of social-orientation

We tested whether MD levels were associated with self- and socially-oriented mental contents in general. Results revealed partial support for our hypotheses: while MD scores were not associated with self-oriented contents (Table S3.9), MD scores were positively associated with a higher degree of social-orientation, *B* = 0.0014, *SE* = 0.0004, *p* < 0.001, *β* = 0.08 (Table S3.10).

To better characterize this social-orientation effect, we examined whether the observed relationships were specific to internally-oriented thinking. As predicted, we found that MD levels significantly moderated the relationship between internal-orientation and social-orientation, such that individuals with higher MD scores showed a stronger positive association between degree of internal-orientation and social-orientation, *B* = 0.0015, *SE* = 0.0005, *p* = 0.007, *β* = 0.02, as shown in (Fig 4; Table S3.11). Said differently, for individuals higher in MD, more internally-oriented thoughts were likely to be more socially-oriented.

**Figure 4.**
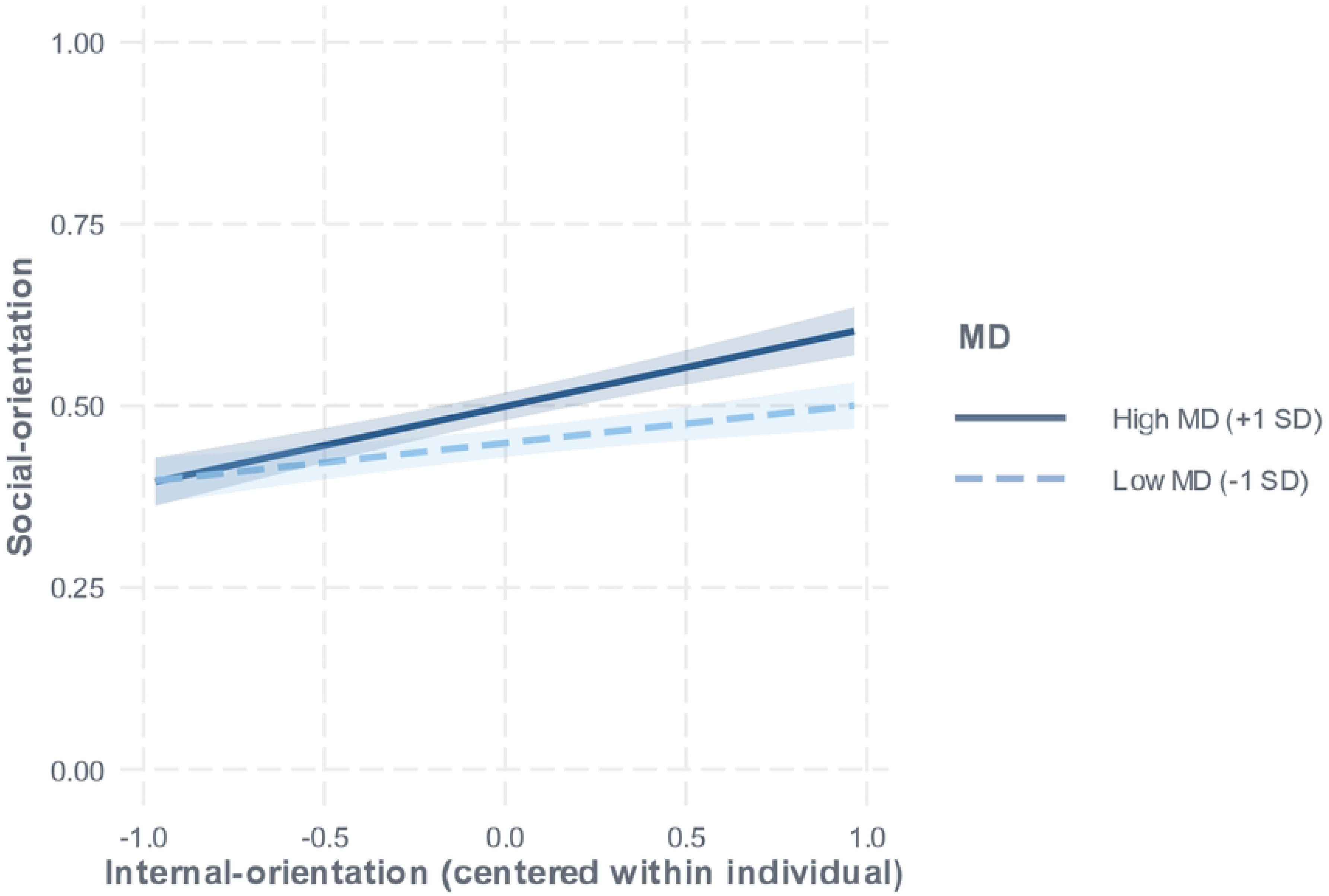
The interaction between the degree of internal-orientation and MD significantly predicted the degree of social-orientation. Simple slopes illustrated the association between internal-orientation and social-orientation at low (−1 SD, dotted line) and high (+1 SD, solid line) levels of MD. Lines indicate fitted regression slopes of the fixed effects, with shading representing 95% CI. Internal-orientation rating was within-subject mean centered.

Finally, we tested whether the association between MD and social-orientation could be explained by differences across and within individuals in social context (being alone or with others). First, higher MD scores were not associated with higher probability of being alone, log odds = 0.0028, odds ratio = 1.003, *SE* = 0.0025, *p* = 0.260, despite positive associations between MD and perceived loneliness (Table S3.12 – S3.14). Next, we examined whether the positive association between MD scores and social-orientation remained significant when restricting analyses to observations in which participants were alone, and indeed found this to be the case (*B* = 0.0017, *SE* = 0.0005, *p* < 0.001, *β* = 0.10). Although being alone was associated with thoughts rated less social-oriented, *B* = -0.1210, *SE* = 0.0051, *p* < 0.001, *β* = -0.38, MD did not significantly moderate this relationship, *B* = 0.0005, *SE* = 0.0003, *p* = 0.061, *β* = 0.03. Thus, associations between MD and greater social orientation does not appear to be explained by differences in social context or simply by being alone.

#### MD was uniquely associated with greater thought persistence

Additional preregistered analyses evaluated the direct and moderating roles of MD symptoms across other momentary thought characteristics (regression tables with the complete set of variables were reported in Supplementary Material S4). Higher MD scores were associated with slightly lower task-relatedness, B = -0.0009, SE = 0.0004, p = 0.014, *β* = -0.06; however, the association was attenuated after controlling for trait rumination, which independently predicted lower task-relatedness *B* = -0.0321, *SE* = 0.0089, *p* < 0.001, *β* = -0.08. Additionally, MD scores were not significantly associated with momentary intentionality B = -0.0004, SE = 0.0004, p = 0.335, *β* = -0.02, while trait rumination was linked to reduced intentionality, B = -0.0429, SE = 0.0107, p < 0.001, *β* = -0.11. In contrast, MD scores were uniquely linked to greater persistence of momentary mental experiences before and after controlling for rumination, B = 0.0011, SE = 0.0003, p < 0.001, *β* = 0.08, while trait rumination showed no unique association with persistence.

Moderation analyses further clarified the distinct cognitive dynamics of MD severity and trait rumination. Contrary to our preregistered hypotheses, we found no evidence that MD levels moderated the relationships between intentionality and persistence, internal- or external-orientation and persistence, intentionality and social-orientation, positive thought content and persistence, or task-relatedness and persistence (Table 5). However, in an independent model evaluating trait rumination alone, rumination negatively moderated the relationship between task-relatedness and thought persistence (B = -0.0262, SE = 0.0091, p_adj_ = 0.025, *β* = -0.02), suggesting that higher levels of rumination were associated with a greater likelihood that more off-task thoughts would be more persistent. Overall, these findings suggest that although MD symptoms are characterized by more persistent and somewhat more task-unrelated mental experiences, it does not appear to be systematically associated with an altered momentary coupling among these cognitive characteristics.

## Discussion

The present study provides new evidence regarding the momentary cognitive and social characteristics associated with maladaptive daydreaming (MD) levels in a large sample utilizing an EMA approach. Overall, MD symptoms were consistently linked to poorer mental health. Higher MD scores were associated with higher degrees of internal-orientation, more negative state affect, more negatively valenced thought content, greater socially-oriented content, and higher persistence of momentary experience. These associations persisted beyond trait rumination, highlighting the unique variance accounted for by MD. By capturing momentary thought dynamics, this study advances understanding of the phenomenology of MD and its role in shaping everyday thought characteristics.

MD symptoms and trait rumination showed a moderate yet reliable association, consistent with the notion that both involve repetitive, internally directed forms of cognition that are difficult to disengage from and associated with psychological distress. Both are characterized by higher automatic constraints. However, the constructs remain conceptually distinct. MD represents immersive narrative imagination that increasingly becomes difficult to regulate [23], whereas rumination represents a maladaptive cognitive process of thinking perseveratively about one’s feelings and problems [44]. The moderate association therefore likely reflects shared self-regulatory vulnerabilities toward excessive internal cognition, while their differences in content and function maintain their conceptual separability.

MD scores remained uniquely associated with lower well-being and poorer mental health even after accounting for rumination. Its negative association with mindfulness suggests that MD is characterized by heightened automatic constraints and reduced regulatory control [45]. This is consistent with earlier evidence that individuals with MD were less attentive to, less able to identify, and less accepting of negative feelings [46] and had lower mindfulness scores [47]. In addition, MD scores were uniquely associated with higher levels of internal-orientation and lower levels of external-orientation, aligning with its definition as habitual engagement in vivid imagined fantasies. In contrast, the association between MD scores and reduced task-relatedness was attenuated when accounting for rumination. Previous research using the Daydreaming Frequency Scale (Polish version) found that daydreaming frequency predicted more daily task-unrelated thoughts and lower daily thought control [48]. The present findings partially extend this literature by showing that MD was similarly associated with lower regulatory control. However, the attenuated relationship after controlling for rumination indicated that these characteristics may not be unique to MD but instead reflect broader patterns of repetitive internal cognition. Thus, lowered regulatory control may be associated with co-occuring rumination rather than being specific to MD. Additionally, MD symptoms were uniquely linked to anxiety and depressive symptoms, consistent with previous studies linking MD to social anxiety [49], generalized anxiety disorder, and depression [50], as well as extensive evidence establishing robust associations between rumination and internalizing symptoms [51,52]. These associations indicate potential shared risk processes such as maladaptive repetitive thinking, through which MD and rumination contribute to psychological distress.

Together, these findings suggest that MD shares several transdiagnostic processes with rumination and internalizing psychopathology, particularly maladaptive repetitive thinking, heightened automatic constraints, and diminished regulatory control over internally generated cognition. At the same time, MD remains phenomenologically distinct in its stronger association with immersive, vivid, and internally focused experiences.

Across the EMA surveys, both MD levels and trait rumination predicted worse state affect and negatively-toned content, each contributing independent variance. This result is consistent with a daily diary study showing that, among individuals with MD, higher levels of depression and more intense daydreaming were linked to greater negative mood [53]. 10/29/2026 9:36:07 AMYet, when our analyses were restricted to surveys only including internally-oriented thoughts, covariation with rumination fully accounted for the negative relationships between MD, negative state affect, and emotional tones of thought contents. Further, a key nuance of the findings was in reconciling the general main effect of MD on lower state affect with the conditional three-way interaction among MD levels, trait rumination, and internal-orientation. When individuals have high co-occurring rumination tendencies, thoughts characterized by greater internal orientation were associated with lower state affect regardless of MD levels. However, when co-occurring trait rumination was low, engaging in internal orientation predicted more positive state affect when MD score was high. Rather than being contradictory, these results suggest that rumination may be the driving factor of negative affect on a momentary basis in explaining the negative relationship between MD scores and state affect. While internally-oriented thoughts in MD episodes may be associated with higher state affect in the moment – perhaps serving an escape or compensatory function to cope with undesirable reality [54] – habitually engaging with such thoughts may have a cumulative emotional cost, particularly when high co-occuring trait rumination overrides the momentary affect benefits.

Interestingly, we found that MD levels had a unique relationship with social cognition. Both MD and trait rumination were independently associated with lower levels of positive relations with others and greater loneliness – two subjective indicators of worse social well-being. These findings are consistent with previous studies showing that rumination was related to poorer perceived interpersonal skills [55] and perceived social support [56]. MD symptoms were also consistently linked to social dysfunction, such as social anxiety disorder [23], loneliness [57], and an ambivalent-fearful attachment style [46]. Despite these negative associations with social well-being, individuals with higher levels of MD exhibited a greater propensity to engage in socially oriented thoughts. This aligns with the account that individuals with MD experience greater social isolation and social impairment and may use MD experience as a coping tool to fulfill interpersonal and emotional needs [4,58]. Brenner and colleagues showed that the majority of respondents endorsed daydreaming about romantic relationships or finding love (81.3%) [58]. Adaptive forms of social daydreaming may be beneficial in satisfying individuals’ need for social connection and improving socio-emotional regulation, especially when thinking about a close other [59]. Similarly, imaginary companions can help satisfy interpersonal needs, cope with loneliness or fear, and improve self-esteem, from childhood to early adulthood [58,60]. However, in the case of MD, socially-oriented thoughts may primarily serve a compensatory function by substituting for unmet real-life interpersonal and emotional needs [60]. Although such fantasies may provide a temporary comfort or a sense of connection, excessive reliance on imagined social interactions may reinforce social withdrawal, highlight discrepancies between desired and actual relationships, and ultimately perpetuate loneliness and negative affect. Details about how people construct their imagined social relationships and who they are daydreaming about in MD episodes are needed to further analyze the mechanism underlying social contents in MD.

Although prior work suggests that social isolation in individuals with MD may provide a space to engage in intensive imaginative experiences [4], contrary to our hypothesis, higher MD scores were not associated with a greater likelihood of being alone. While individuals with MD may isolate themselves to immerse in daydreaming or avoid judgement or interruptions [12], our data only represents a sample of daily thoughts rather than specific MD episodes, which may account for this null finding.

Contrary to our preregistered hypotheses, MD levels did not moderate any within-person association among momentary thought characteristics, including the relationships between intentionality and persistence, internal orientation and persistence, external orientation and persistence, or task-relatedness and persistence. MD severity was linked to elevated thought persistence regardless of the specific type or content of the thought, suggesting a more general tendency towards longer and stickier thoughts and attention across the board. On the contrary, trait rumination did not demonstrate main effect relationships with thought persistence, yet was associated with lower overall task-relatedness and an interaction such that at higher levels of trait rumination, more task-unrelated thoughts tended to be more persistent. These findings suggest that trait rumination may selectively disrupt the persistence of task-related thought in MD.

This pattern is consistent with Burrel and colleagues’ [11] extensions to the Dynamic Framework of Thought [12], suggesting that MD manifests as a global elevation in automatic constraints regardless of deliberate constraint. In the present study, MD severity was unrelated to intentionality, and its association with reduced task-relatedness was fully accounted for by trait rumination, with a robust and unique main effect of MD on thought persistence. The thought persistence may reflect the compulsiveness and stickiness compoments of MD, which is not modulated by the presence of deliberate constraints: fantasy can be initiated either spontaneously (unprompted by an affective trigger) or deliberately (initiated with intermittent cognitive control guiding the unfolding of fantasy), while the persistence was experienced across thoughts, including fantasies presumably. Trait rumination, in the Dynamic Framework of Thought, exihibited a different profile marked by impaired or reduced deliberate constraint along with heightened automatic constraint. Whereas task-related thoughts naturally sustain attention in healthy cognition, higher trait rumination weakens this association, causing task-focused mental activity to lose its persistence and become easily disrupted, which easily and repeatedly draw attention back to repetitive negative content. Although our findings are somewhat speculative and should be extended with experimental manipulations, our finding suggests that the “stickiness” of rumination may differ qualitatively from that of MD.

The use of EMA represents a methodological strength of this study, as it enables the capture of real-time, ecologically valid thought characteristics. This is particularly important because this is the first EMA study to our knowledge to examine MD-related momentary cognitive characteristics in everyday life. Rather than identifying discrete MD episodes in real time, this study assessed how individuals with varying levels of MD differ in their overall daily cognitive profiles. This approach provides valuable insight into trait-level differences in thought patterns; however, it also limits our understanding of the immediate features, antecedents, and consequences of MD episodes. Our data may not fully reflect the richness or variability of the complete cognitive profile due to participant compliance. Future studies may adopt event-contingent sampling to specifically study the cognitive characteristics of MD episodes among individuals with MD.

Trait rumination was included in this study as a comparison construct and a control variable to test the independent effect of MD beyond general repetitive negative thinking tendencies.

Consistent with this framing, previous literature has noted that MD mechanisms share similarities with obsessive thinking and compulsion, as all are characterized by intrusive and difficult-to-control features [7,23]. Thus, this overlap suggests that future research should further incorporate obsessive–compulsive symptoms to clarify the boundaries of MD and better situate it within the broader landscape of maladaptive repetitive thought. Overall, these findings suggest that MD represents a distinctive form of maladaptive internally directed cognition characterized by immersive imaginative experiences and social fantasy, while simultaneously sharing transdiagnostic self-regulatory vulnerabilities with rumination and potentially other forms of repetitive thought. By characterizing the everyday cognitive and affective correlates of MD, the present study provides a foundation for future work aimed at clarifying its mechanisms and informing targeted interventions.

## Acknowledgement

The authors would like to thank Matthew Grilli for input on the Mind Window app used for this project, and Delaney Freveletti and Freya Abraham for assistance with data collection.

ChatGPT (Open AI) and Gemini (Google) were used to assist with searching relavent literature, debugging R code, summarizing synthesized results, and refining language for grammar, clarity, and readability. No identifiable data or raw data was shared with any AI tools. The authors have carefully reviewed and edited all AI output and take full responsibility for the content.

